# Alternating Polarity as a Novel Strategy for Cultivating Electro-Methanogenic Microbial Communities Capable of Robust Biogas Production

**DOI:** 10.1101/2023.08.29.555408

**Authors:** Shiyun Yao, Clifford S Swanson, Zhang Cheng, Qiang He, Heyang Yuan

## Abstract

Electro-methanogenic microbial communities can produce biogas with high efficiency. Extensive efforts have been made to cultivate these communities in engineered systems. Conventional cultivation strategies can select electrotrophic methanogens but not their electron-donating partners, resulting in communities that are sensitive to perturbations. Herein, we developed an alternating polarity strategy to simultaneously select both microbial populations. In two-chamber bioelectrochemical systems amended with activated carbon, the electrode potential was alternated between +0.8 V and -0.4 V vs. standard hydrogen electrode every three days. After eight alternating cycles, cultivated activated carbon was transferred into new bioreactors, and the enrichment procedure was repeated four times. Cumulative biogas production under alternating polarity increased from 45 L/L/kg-activated carbon after start-up to 125 L/L/kg after the 4^th^ enrichment, significantly higher than that under intermittent cathode (-0.4 V/open circuit), continuous cathode (-0.4 V), and open circuit. The communities cultivated under alternating polarity were electroactive and structurally different from those cultivated under other conditions. One *Methanobacterium* population and two *Geobacter* populations were consistently abundant and active in the communities. Their 16S rRNA was upregulated by electrode potentials. Bayesian networks inferred close associations between these populations. The cultivation strategy can enhance biogas production, and the cultivated communities may serve as a model system for elucidating the mechanisms of extracellular electron uptake.

**Synopsis:** An alternating polarity strategy was developed in this study to cultivate electro-methanogenic microbial communities. The cultivated communities can produce biogas more efficiently and help us understand the ecophysiology of the key microbial populations.

## 1. Introduction

Methanogenic microbial communities are cultivated in engineered systems to convert organic wastes such as waste activated sludge and food waste into methane-rich biogas [1, 2], a renewable energy source that has attracted considerable attention [3-5]. Biogas is produced cooperatively by several groups of functional microbial populations in the communities [6, 7]. These include primary fermentative bacteria that degrade complex organic matter to monomers, acetogens that produce acetate, syntrophs that produce hydrogen, and methanogens that scavenge these fermentation products for acetoclastic and hydrogenotrophic methanogenesis [8, 9]. The availability of fermentation products can strongly affect methanogenic activity and represents one of the limiting factors for biogas production [9-13].

Recent studies show that biogas production can be enhanced by cultivating electro-methanogenic communities [14, 15]. In addition to common fermentative bacteria, electro-methanogenic communities consist of electrotrophic methanogens and electroactive bacteria (primarily *Geobacter*) [16]. Electrotrophic methanogens can use cathodes as electron donors to reduce carbon dioxide to methane [17-19]. Alternatively, they can accept extracellular electrons from electroactive *Geobacter* [20-22], a syntrophic relationship also known as direct interspecies electron transfer (DIET) [23-25]. Compared to acetoclastic and hydrogenotrophic methanogens, the activity of electrotrophic methanogens is not limited by the availability and diffusion of acetate and hydrogen [26-28]. As a result, systems harboring electro-methanogenic communities can produce biogas at a higher rate [29-32]. There is continued interest in developing strategies to cultivate electro-methanogenic communities.

Electro-methanogenic communities can be assembled using either bottom-up or top-down strategies [33]. Bottom-up strategies, i.e., coculturing electrotrophic methanogens and electroactive bacteria [34-36], are used to understand their physiology but are not suitable for practical applications. Top-down strategies, on the other hand, can be used to cultivate communities with a minimum number of functional populations by imposing selective pressure [33]. This is particularly attractive for biogas production systems where the community structure is continuously influenced by microbial immigration from the feed substrate [37-39]. Common top-down strategies for cultivating electro-methanogenic communities include cathode enrichment, microbial electrolysis, and conductive material amendment. Cathode enrichment refers to using a cathode as an electron donor to select electrotrophic methanogens. The cathodic potential is typically between -0.24 and -0.41 V vs. standard hydrogen electrode (SHE) to avoid hydrogen evolution and selection of hydrogenotrophic methanogens [40]. Similar to cathode enrichment, microbial electrolysis is an electrochemical process in which an external voltage is applied to drive electron transfer from the anode to the cathode [41]. Electroactive bacteria and electrotrophic methanogens are selected in the anode and cathode chambers, respectively [42]. In contrast, conductive material amendment does not impose electrochemical stimulation, but the materials act as a conduit that promotes extracellular electron transfer from electroactive bacteria to electrotrophic methanogens [43].

These top-down strategies focus on the selection of electrotrophic methanogens but overlook the simultaneous selection of electroactive bacteria. Without electroactive bacteria acting as an alternative electron donor, the biogas production performance of the communities can be compromised by perturbations [44]. For example, during cathode enrichment, the electrode is the sole electron donor, and disconnecting the electrode for a short period of time (e.g., 45 min) can result in a significant, irreversible decrease in biogas production rate [40]. During microbial electrolysis, the electroactive bacteria selected on the anode are unable to interact with the electrotrophic methanogens on the cathode because the distance between the two electrodes is orders of magnitude greater than the thickness of the electroactive biofilm (e.g., 10 cm vs. 10 μm) [42]. In addition, the cathodic potential during microbial electrolysis is not precisely controlled and is often more negative than -0.41 V vs. SHE (redox potential of hydrogen evolution at pH 7) [45], leading to the selection of hydrogenotrophic methanogens [41, 46]. Finally, conductive materials do not create a strong driving force for cultivating electro-methanogenic communities [14]. In systems amended with conductive materials, *Geobacter*, which is by far the only confirmed electron-donating partner involved in DIET [15], can occur with high abundance [28, 47, 48]. However, their dominance does not necessarily indicate electrotrophic methanogenesis [14], as *Geobacter* can transfer electrons extracellularly to other electroactive bacteria or grow syntrophically with hydrogenotrophic methanogens [31, 49-51].

We hypothesize that electro-methanogenic communities can be effectively cultivated with alternating polarity (Figure 1). Alternating polarity refers to switching the electrode potential periodically between cathodic and anodic potentials [52, 53]. When the potential is alternated, the electrode acts as a stable electron donor and acceptor for electrotrophic methanogens and electroactive bacteria, respectively. Consequently, the two populations are simultaneously selected in a microenvironment (e.g., on the surface of conductive materials) and can interact with each other for robust electrotrophic methanogenesis. Our hypothesis is supported by an early study, in which a bioelectrochemical system achieved faster methane production when the electrode periodically served as the anode and cathode [54]. One-time reversal of the electrode potential has also been reported to promote the formation of electroactive biofilm, shorten reactor start-up, and enhance methane production [54-56]. It should be noted that the electrode potentials in those studies were not precisely controlled, which could lead to the selection of undesirable populations [41, 46]. Moreover, the alternation frequency needs to be carefully controlled to allow the microbes, in particular electrotrophic methanogens, to adapt to the changing potentials. High alternation frequency can cause permanent damage to the electroactive biofilm [57], and the resulting communities are incapable of methanogenesis [58-61].

**Figure 1.**
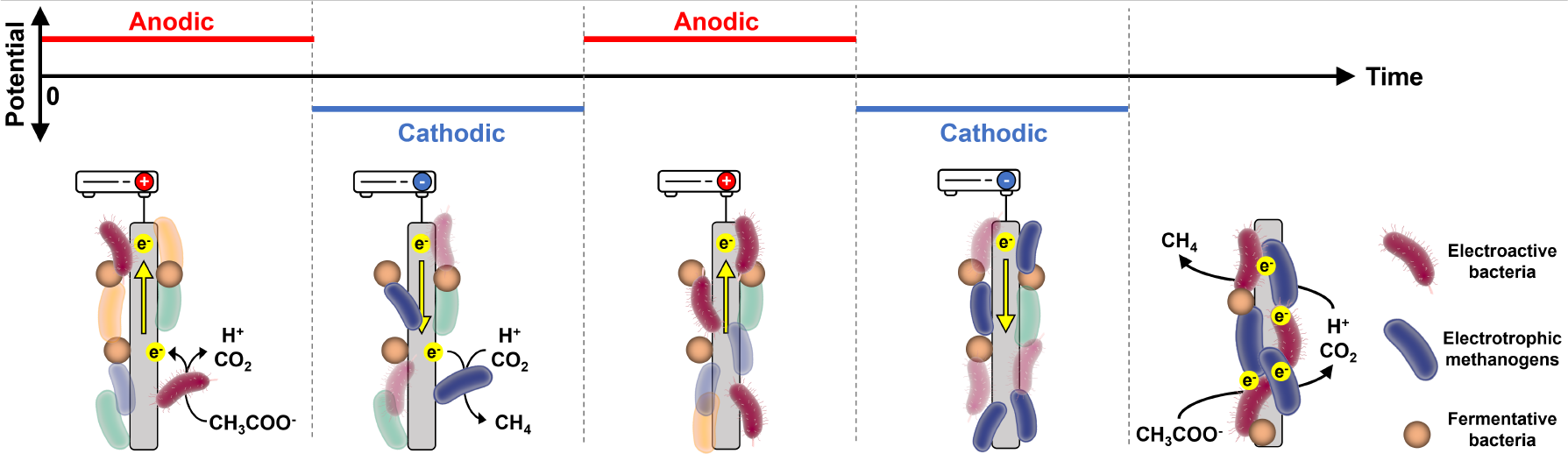
Schematic of cultivating electro-methanogenic communities using alternating polarity.

The goal of this study is to develop an alternating polarity strategy to cultivate electro-methanogenic communities capable of robust biogas production. To this end, two-chamber bioelectrochemical systems were amended with granular activated carbon (a conductive material), and the potential of the working electrode was precisely controlled using a potentiostat. The cathodic potential was set at -0.4 V vs. SHE for electrotrophic methanogens to use as an electron donor without selecting hydrogenotrophic methanogens. An anodic potential of +0.8 V vs. SHE was used for electroactive bacteria to use as an electron acceptor. Additionally, an open-circuit cycle was performed between the cathodic and anodic cycles to promote the interaction between the two populations [62]. Each condition lasted for three days to mitigate possible electrochemical shock. Bioelectrochemical systems operated under intermittent cathode (-0.4 V vs. SHE and open circuit), continuous cathode (-0.4 V vs. SHE), and open circuit were used as control groups. The microbial communities were characterized with 16S rRNA gene amplicon sequencing, and the activity of the microbial populations was analyzed with 16S rRNA transcript profiling and Bayesian networks. To our best knowledge, this is the first attempt to use alternating polarity to cultivate electro-methanogenic communities. The strategy may hold great promise in practical applications, and the communities may be used as a model system to reveal the different mechanisms by which electrotrophic methanogens perform extracellular electron uptake from cathodes and electroactive bacteria [16].

## 2. Materials and Methods

### 2.1 Experimental design

Two-chamber bioelectrochemical systems were constructed as previously described [51]. Briefly, the methanogenic chamber was amended with 50 g of granular activated carbon, resulting in a liquid volume of 60 mL. A carbon brush was buried in activated carbon as the working electrode, and an Ag/AgCl reference electrode (+0.197 V vs. SHE) was installed adjacent to the working electrode. The methanogenic chamber was inoculated with digester sludge supernatant collected from a local wastewater treatment plant and fed with synthetic soft drink wastewater [63]. The synthetic wastewater contained 3,000 mg/L fructose and 2,200 mg/L polyethylene glycol 200 as carbon sources and was slightly modified with 0.1 M phosphate buffer saline to maintain the pH. Stainless steel mesh was used as the counter electrode in another chamber. The counter-electrode chamber was filled with 0.1 M phosphate buffer saline. The two chambers were separated using a cation exchange membrane (Membrane International Inc.), and the solutions in both chambers were recirculated using a peristaltic pump at a rate of 60 RPM. The schematic of the bioreactor can be found in Supporting Information (SI) Figure S1.

Eight bioreactors were constructed and operated in batch mode with a hydraulic retention time of three days. Duplicate bioreactors were operated under one of four cultivation conditions (Table 1): 1) the potential was alternated between +0.8 and -0.4 V vs. SHE, 2) the potential was intermittently set at -0.4 V vs. SHE, 3) the potential was continuously set at -0.4 V vs. SHE, and 4) no potential was applied (open circuit). The electrode potential was poised using a multi-channel potentiostat (Squidstat™ Prime, Admiral Instruments), and each cultivation condition was examined for eight cycles. The duration of each cycle was the same as the hydraulic retention time (3 days). After eight cycles, start-up was completed, and 10% (w/w) of the cultivated activated carbon was transferred to new bioreactors containing fresh activated carbon. The newly inoculated bioreactors were again operated under the same cultivation conditions as in prior cycles for eight more cycles. Enrichment was repeated four times. The schematic of the enrichment procedure can be found in SI Figure S1.

**Table 1.** Electrode potentials applied in the eight cycles in one enrichment. Each cycle lasted for three days.

| Condition | Electrode potential (V vs. SHE) |  |  |  |  |  |  |  |
| --- | --- | --- | --- | --- | --- | --- | --- | --- |
|  | Cycle 1 | Cycle 2 | Cycle 3 | Cycle 4 | Cycle 5 | Cycle 6 | Cycle 7 | Cycle 8 |
| Alternating | +0.8 | - | -0.4 | - | +0.8 | - | -0.4 | - |
| Intermittent | -0.4 | - | -0.4 | - | -0.4 | - | -0.4 | - |
| Continuous | -0.4 | -0.4 | -0.4 | -0.4 | -0.4 | -0.4 | -0.4 | -0.4 |
| Open circuit | - | - | - | - | - | - | - | - |

### 2.2 Chemical and electrochemical analyses

Biogas were collected at the end of each cycle using gas sampling bags, and the volume was measured manually using syringes and normalized to the volume of the substrate and the mass of activated carbon (L/L/kg-activated carbon). Methane content was measured using gas chromatography–mass spectrometry (8860/5977B, Agilent Co.) equipped with an Agilent CP-Molsieve 5A column (25 m X 0.25 mm) according to an established method [64]. Effluent was collected after each cycle. The pH and soluble chemical oxygen demand (COD) in the effluent were measured using a pH meter (Mettler-Toledo LLC.) and a standard colorimetric method, respectively. Electric current was recorded using the built-in software of the potentiostat. After the last cycle of the final enrichment, cyclic voltammetry (CV) was conducted to characterize the electroactivity of the communities. Voltammograms were obtained with fresh substrate (turnover condition) at different scan rates (0.2, 1, 2, 5, 10, 20, and 50 mV/s) and a scan window of -0.4 V to +0.8 V vs. SHE, which covered the applied potentials and the redox potentials of most electroactive microbes.

### 2.3 Biomass collection and sequencing

Biomass samples (1.5 g of cultivated activated carbon) were collected after Cycles 7 and 8 in each enrichment. Additional biomass samples were collected from the alternating-polarity bioreactors after Cycles 5 and 6 in the 1^st^ and 4^th^ enrichments. Samples were stored at -80 °C prior to nucleic acid extraction. Additionally, samples for RNA extraction were stored in a RNAlater stabilization solution. Genomic DNA was extracted using the MPbio FastDNA Spin kit (MP Biomedicals). RNA was extracted using the QIAGEN RNeasy PowerSoil Total RNA kit (Qiagen) followed by removal of residual DNA using the QIAGEN Dnase Max DNA-free kit (Qiagen) and reverse transcription using the SuperScript IV VILO Master Mix (Thermo Fisher Scientific). The DNA and cDNA samples were quality-checked and quantified using Nanodrop and the Thermo Fisher Qubit™ dsDNA Quantification Assay (Thermo Fisher Scientific).

After DNA extraction, the V4 region of the 16S rRNA gene and rRNA were amplified using polymerase chain reaction (PCR) with the primer pair 515F (GTGCCAGCMGCCGCGGTAA) and 806R (GGACTACHVGGGTWTCTAAT, with unique barcodes) [65]. The amplicons were quality-checked using gel electrophoresis, and amplicons with a length of 300 bp were targeted. Subsequently, the amplicons were purified with the ChargeSwitch Nucleic Acid Purification Technology (Invitrogen) and then pooled to equimolar based on the quantification results with NanoDrop ND-3300 fluorospectrometer (Thermo Fisher Scientific). The pooled library was further quantified with the KAPA Illumina Library Quantification Kit (Kapa Biosystems) and finally sequenced on an Illumina MiSeq platform (Illumina). Paired-end sequences were assembled and denoised using QIIME 2, and operational taxonomic units (OTUs) were picked using DADA2 [66, 67]. Taxonomy was assigned using the QIIME 2 plugin feature-classifier and the SILVA database as the reference, with the classifier trained on 99% OTUs [68, 69]. Raw sequencing reads were deposited in the National Center for Biotechnology Information database under accession number PRJNA1004309.

### 2.4 Microbial community analysis

After singletons were removed, statistical analyses were performed using the software R. These included two-sample t-test, analysis of variance (ANOVA), and Bray-Curtis dissimilarity-based principal coordinate analysis (PCoA). PCoA were performed with relative abundance (16S rRNA gene). Permutational multivariate analysis of variance (PERMANOVA) was carried out to test the significant difference of the PCoA results (N = 999). A *p*-value < 0.05 was used to identify a significant difference. Core populations were selected at the OTU level based on the following criteria: average relative abundance > 0.75% and occurrence > 75% across all DNA samples. A phylogenetic tree of the core populations was constructed using the software ARB and a neighbor-joining method [70].

Bayesian networks were constructed using the R package “bnlearn” to infer the interactions of the microbial populations [71]. Briefly, the relative abundance of 16S rRNA of the core OTUs and three key environmental factors (biogas production, cathodic current, and pH) were combined as a single dataset and normalized to between 0 and 1:

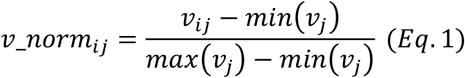

where *v_norm*_*ij*_ is the normalized variable *j* in sample *i, v*_*ij*_ is the observed variable *j* in sample *i, min(v*_*j*_) and *max(v*_*j*_) are the minimum and maximum values of variable *j*. The network structure was learned using a hill-climbing algorithm, and the network parameters were learned using a maximum likelihood method [72]. The networks were tested with leave-one-out cross validation [73]. In each cross-validation cycle, the relative root-mean square error (RMSE) between the observed and predicted values was calculated:

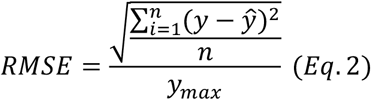

where *y* is the experimental value, _ŷ_ is the predicted value, and *y*_*max*_ is the maximum experimental value. Null model predictions were performed by manually setting the value of the node of interest (e.g., the abundance of a population or the measurement of an environmental factor) to the mean value across all samples.

## 3. Results and Discussion

### 3.1 Activity of the electro-methanogenic communities

The overall activity of the microbial communities cultivated with alternating polarity (+0.8/-0.4 V), intermittent cathode (-0.4 V/open circuit), continuous cathode (-0.4 V), and open circuit was characterized with bioreactor performance including biogas production, methane content, COD removal, effluent pH, and voltammetric responses. Biogas production as a key indicator of a functioning methanogenic community was expressed as cumulative biogas produced per unit volume of substrate and unit mass of activated carbon (Figure 2A). At the end of the start-up period, alternating polarity resulted in 45 L/L/kg of biogas, which was significantly lower than the 57 L/L/kg achieved with continuous cathode (two-sample *t*-test, *p* < 0.05) but comparable to that obtained with intermittent cathode and open circuit. After eight cycles of cultivation, 10% of the activated carbon was used to inoculate new bioreactors containing fresh activated carbon. This enrichment procedure enhanced biogas production. The biogas produced under alternating polarity increased from 56 L/L/kg after the 1^st^ enrichment to 65 L/L/kg and 80 L/L/kg after the 2^nd^ and 3^rd^ enrichments, respectively. At the end of the 4^th^ enrichment, alternating polarity yielded 125 L/L/kg biogas and outperformed all other conditions, indicating the formation of electro-methanogenic communities that were highly efficient in biogas production.

**Figure 2.**
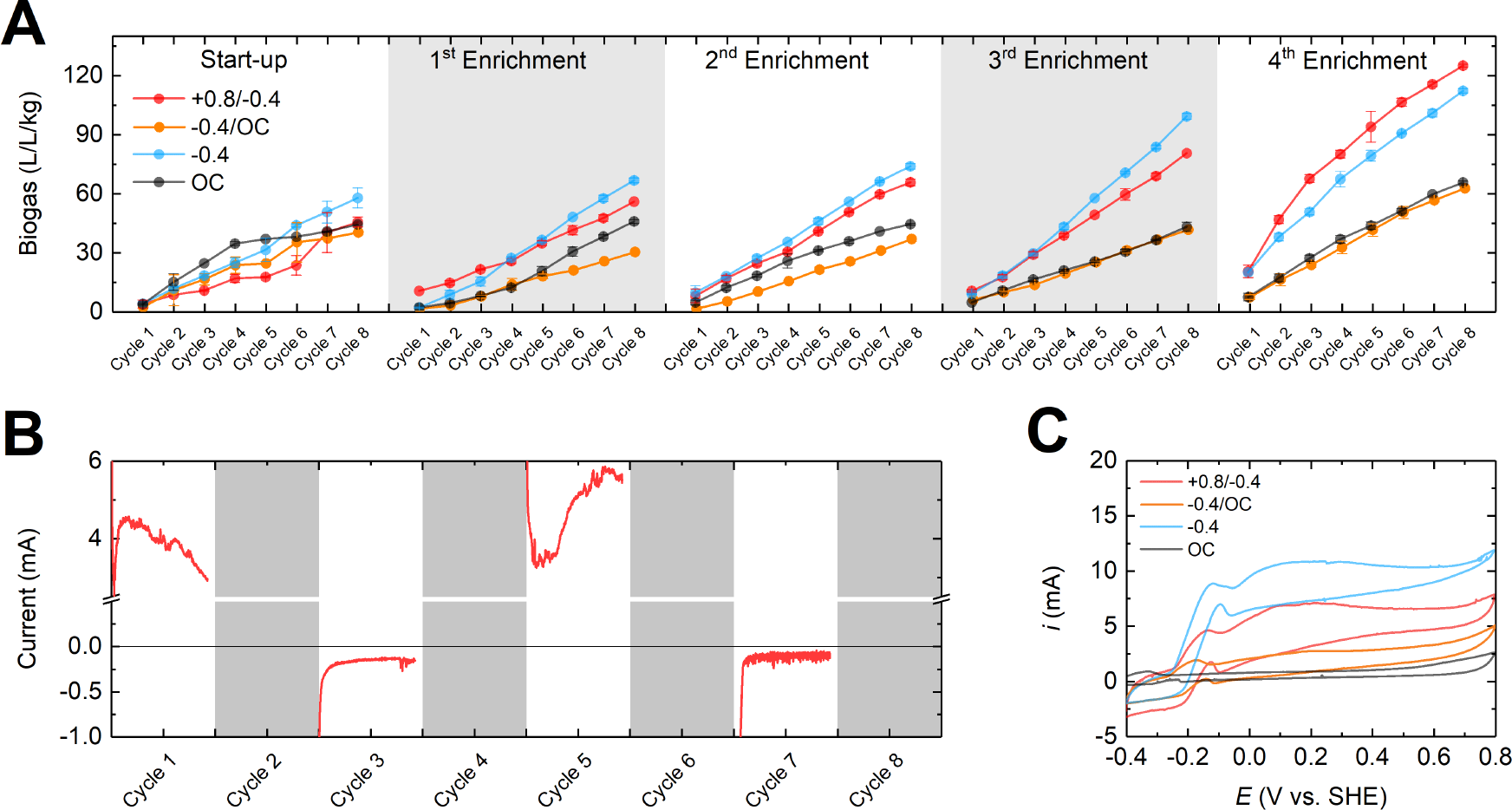
(A) Cumulative biogas production during the start-up and four enrichments. (B) Current production by the alternating polarity-cultivated communities in the 4^th^ enrichment. (C) CV conducted with a scan rate of 0.2 mV/s.

The average methane content (∼66%) and COD removal (∼3,500 mg/L) did not differ significantly under different cultivation conditions (SI Figure S2). The effluent pH, on the other hand, varied in response to the electrode potential. While the effluent from the intermittent-cathode, continuous-cathode, and open-circuit bioreactors remained neutral (∼7.5) throughout the cultivation, the effluent pH under alternating polarity repeatedly dropped below 6 in the first and fifth cycles when +0.8 V was applied (SI Figure S3). Anode acidification occurs in bioelectrochemical systems as electroactive bacteria use the anode as an extracellular electron acceptor to oxidize organic matter into carbon dioxide and proton [74]. In line with the decreased pH, the alternating-polarity bioreactors produced approximately 4 mA of anodic current (Figure 2B). This led to an average Coulombic efficiency of 54% in the anodic cycles, which was much higher than the 5% in our previous study and indicated the high activity of electroactive bacteria [51]. The cathodic current of the alternating-polarity and intermittent-cathode bioreactors was around -0.2 mA (Figure 2B and SI Figure S4), whereas that of the continuous-cathode bioreactors reached up to -10 mA (SI Figure S4). The higher efficiency of the continuous cathode-cultivated communities in accepting extracellular electrons from the cathode could explain their higher biogas production rate in the first three enrichments (Figure 2A).

To further evaluate the electroactivity of the communities, CV was performed at a low scan rate of 0.2 mV/s (Figure 2C). Typical sigmoidal voltammograms with peak currents were obtained with the alternating-polarity, intermittent-cathode and continuous-cathode bioreactors, suggesting the presence of redox species. In contrast, no peak currents were observed in the voltammogram of the open-circuit bioreactors. Using the first derivative of the voltammograms, the redox potentials of the species were estimated to be -0.2 V and 0 V vs. SHE (SI Figure S5) [75]. These were more positive than the redox potential of carbon dioxide reduction to methane at pH 7 (-0.24 V vs. SHE) and were consistent with typical redox potentials of electroactive *Geobacter* [76, 77]. CV was further performed with the scan rate ranging from 1 to 50 mV/s. The peak current in the voltammograms was found to be strongly correlated with the square root of the scan rate (R^2^ > 0.98, SI Figure S5), implying a reversible and diffusion-controlled electron transfer process [78]. The results collectively confirmed the electroactivity of the electro-methanogenic communities cultivated with electrode potentials.

### 3.2 Dynamics of the electro-methanogenic communities

The effects of electrode potential and enrichment on microbial community dynamics were studied using Bray-Curtis dissimilarity-based PCoA. As shown in Figure 3A, communities formed under different cultivation conditions still overlapped after start-up. This indicates that electrode potential alone may not act as a strong driving force for microbial community assembly. However, combining electrode potential and enrichment significantly changed the community structure. For example, the communities from alternating polarity started to shift since the 1^st^ enrichment and reached the lower left corner after the 4^th^ enrichment (Figure 3B). Communities cultivated under other conditions also shifted but toward different directions (Figures 3C-3E), resulting in clusters that were clearly separated from those in the alternating-polarity bioreactors at the end of the 4^th^ enrichment (Figure 3A). Overall, the results show successful cultivation of distinct electro-methanogenic communities with alternating polarity.

**Figure 3.**
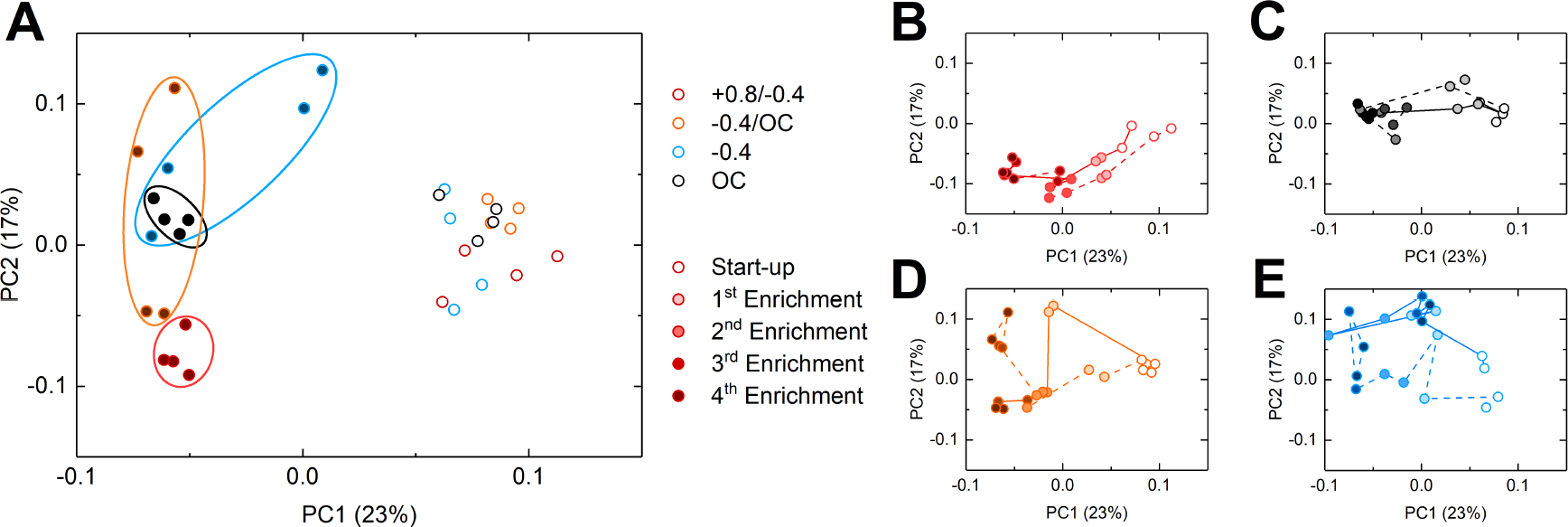
Bray-Curtis dissimilarity-based PCoA. (A) Difference in community strucutre after start-up and 4^th^ enrichment. Community dynamics under (B) alternating polarity, (C) open circuit, (D) intermittent, and (E) cathode continuous cathode. Darker colors indicate later enrichments.

A detailed examination of community dynamics during enrichment suggests that alternating polarity may be a more reliable approach for cultivating electro-methanogenic communities. In particular, the communities in the duplicate alternating-polarity bioreactors shifted in a synchronized manner as the enrichment proceeded (Figure 3B). The communities in the duplicate open-circuit bioreactors also clustered closely after each enrichment (Figure 3C), which was expected as the enrichment procedure did not represent a strong perturbation. In contrast, intermittent and continuous cathodes caused noticeable divergence in community structure. The two communities in the duplicate intermittent-cathode bioreactors evolved along different trajectories and were distant from each other after the 4^th^ enrichment (Figure 3D). A similar differentiation was observed for the communities in the continuous-cathode bioreactors (Figure 3E). The difference in community dynamics under different conditions highlights the important role of the anodic potential (+0.8 V) in cultivating stable electro-methanogenic communities. This may be achieved through the selection of electroactive bacteria, which serve as alternative electron-donating partners and help maintain the community structure under perturbations.

### 3.3 Core populations in the electro-methanogenic communities

OTUs with an average relative abundance greater than 0.75% and occurred in more than 75% of the DNA samples were defined as core populations [79]. Based on this criterion, 22 core populations were selected (Figure 4), accounting for 63% of the total relative abundance. The core populations could be further divided into three guilds potentially responsible for distinct functions: methanogens, *Geobacter*, and fermentative bacteria.

**Figure 4.**
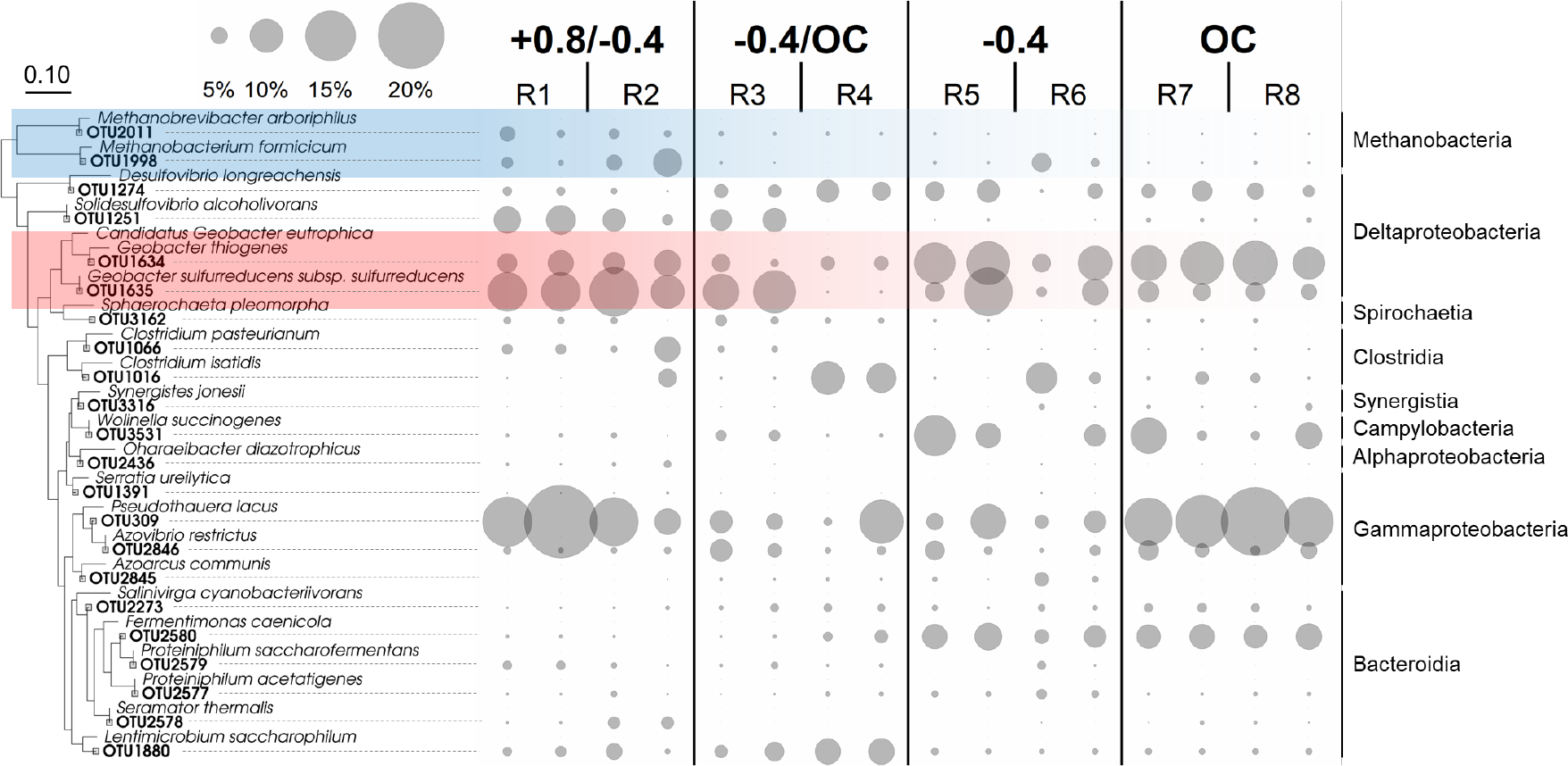
Phylogenetic tree and relative abundance of the 22 core OTUs in the 4^th^ enrichment. The four samples under each cultivation condition were collected from duplicate bioreactors after Cycles 7 and 8.

Two methanogens were identified as core populations: OTU1998 and OTU2011. OTU1998 was phylogenetically close to *Methanobacterium formicicum* and had a high relative abundance of 5% - 10% during start-up and early enrichments in all bioreactors (SI Figure S6). As the enrichment proceeded, its abundance was maintained at about 5% under alternating polarity but dropped below 1% under other cultivation conditions (ANOVA, *p* < 0.05) (Figure 4). This result was consistent with the findings of our previous study [51], in which we applied anodic potentials (0 V and +0.4 V vs. SHE) to select electroactive bacteria but observed high abundances of three *Methanobacterium*-related populations. Several *Methanobacterium* spp. have been observed as dominant methanogens in bioelectrochemical systems and have been speculated to perform extracellular electron uptake from both electroactive bacteria and cathodes [22, 80]. The higher abundance of OTU1998 in the presence of anodic potentials than at intermittent and continuous cathodic potentials demonstrates its ability to accept electrons from anode-selected electroactive bacteria. The other methanogen, OTU2011, was from the genus *Methanobrevibacter*. Its relative abundance remained stable at about 1% under different cultivation conditions (SI Figure S6) and was slightly higher in the alternating-polarity bioreactors than in other bioreactors in the 4^th^ enrichment (ANOVA, *p* < 0.05) (Figure 4). *Methanobrevibacter* spp. are hydrogenotrophic and ubiquitous in various environments [81], but their electrotrophic activity has not been documented.

Two *Geobacter*-related OTUs, OTU1634 and OTU1635, were among the dominant populations throughout the enrichment (SI Figure S6). The selection of OTU1634 appeared to be less efficient with electrode potentials than under open circuit. In the 4^th^ enrichment, the relative abundance of OTU1634 was 4-7% in the presence of anodic and/or cathodic potentials but reached 12% in the open-circuit bioreactors, making it the second most abundant population (Figure 4). OTU1634 shares 97% similarity with *Candidatus* Geobacter *eutrophica*, a novel *Geobacter* species discovered in our previous study [82]. Electrochemical stimulation and omics analysis revealed *Ca*. G. *eutrophica*’s capability of extracellular electron transfer to electrodes and methanogens [51]. *Geobacter*-related OTU1634 may carry similar metabolic pathways, and its lower abundance under alternating polarity implies its low growth efficiency with the anode as an electron acceptor. In contrast, the anode favored the selection of the other *Geobacter*, OTU1635. Its abundance was consistently higher than 8% under alternating polarity (SI Figure S6) and increased to 12% in the 4^th^ enrichment (Figure 4), significantly higher than the 6% under other conditions (ANOVA, *p* < 0.05). With 100% similarity to *Geobacter sulfurreducens*, a model species for extracellular electron transfer [83, 84], OTU1635 was very likely electroactive. However, *G. sulfurreducens* is not known to act as an electron-donating partner in DIET, and therefore the involvement of *Geobacter*-related OTU1635 in electrotrophic methanogenesis remains an open question.

In addition to methanogens and *Geobacter*, the communities cultivated with alternating polarity were also dominated by three fermentative bacteria from the families Clostridiaceae, Desulfovibrionaceae, and Rhodocyclaceae. Among them, *Clostridium*-related OTU1066 was highly abundant (18%) in the 1^st^ enrichment (SI Figure S6). Based on its high similarity to *Clostridium pasteurianum* (100%), OTU1066 was likely responsible for converting carbon hydrates to short-chain volatile fatty acids, ethanol, and hydrogen [85]. This population was gradually replaced by *Solidesulfovibrio*-related OTU1251 and *Pseudothauera*-related OTU309, whose abundances in the 4^th^ enrichment were 7% and 15%, respectively (Figure 4). In comparison, the communities cultivated with intermittent and continuous cathodes showed a much less defined structure without significantly dominant populations (i.e., many populations with low abundances) (SI Figure S6).

### 3.4 Activity and interactions of the core populations

The activity of the core populations was characterized with 16S rRNA transcript profiling. With an average transcript abundance of 3% across all samples, *Methanobacterium*-related OTU1998 was identified as one of the most active populations (SI Figure S7). Its activity was significantly stimulated by alternating polarity than by other conditions (ANOVA, *p* < 0.05). Under alternating polarity, OTU1998 became progressively more active as the enrichment proceeded (Figure 5A), and its 16S rRNA expression was consistently upregulated when electrode potentials were applied (Figure 5B). For example, during start-up, its average transcript abundance was 3%, and the transcript abundance at anodic and cathodic potentials was 0.2 and 1.4 folds relative to that at open circuit, respectively. These numbers increased to 9%, 0.5 folds, and 3.1 folds in the 4^th^ enrichment.

**Figure 5.**
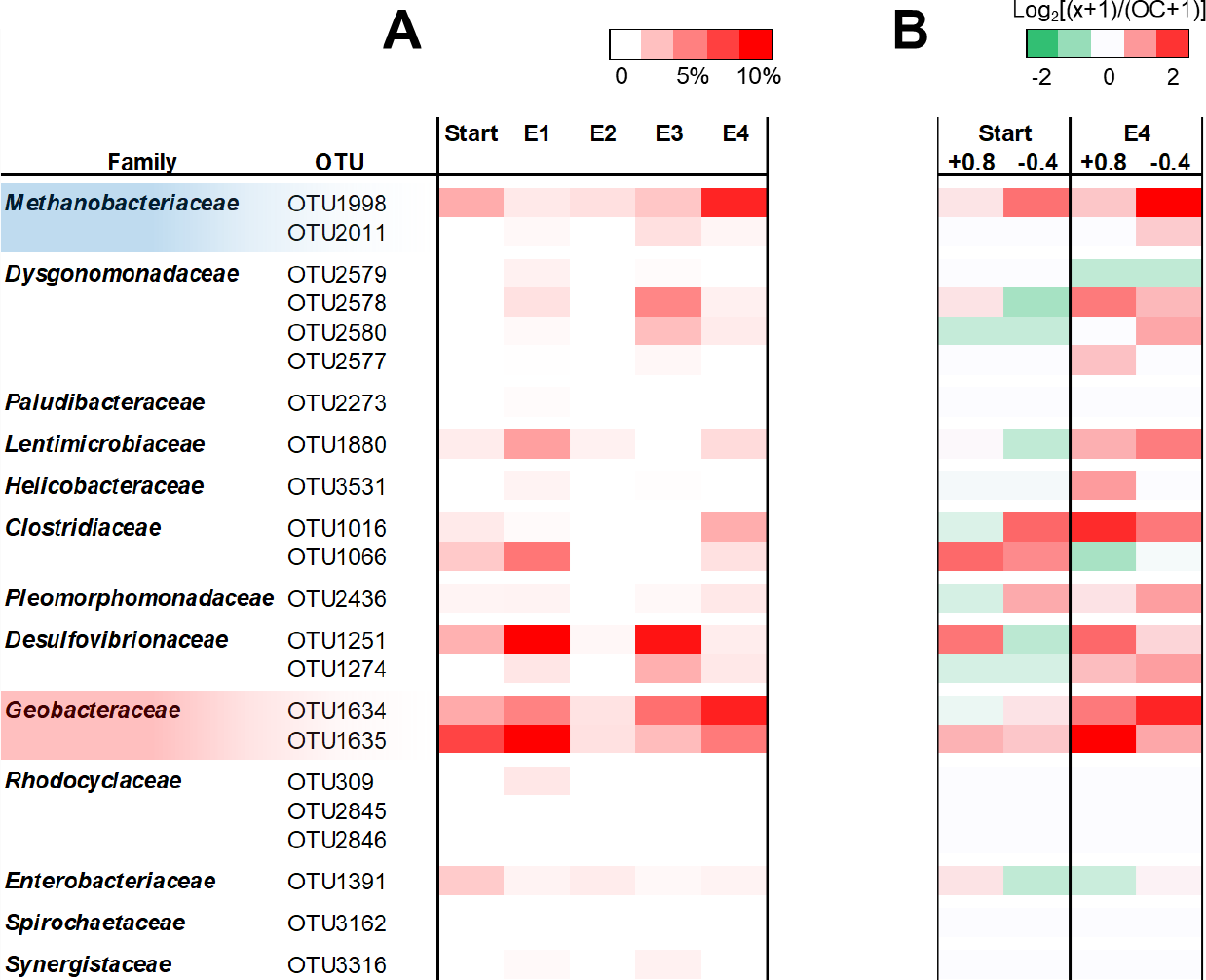
(A) Activity (relative abundance of 16S rRNA transcript) of the 22 core populations under alternating polarity. (B) Change in activity at anodic and cathodic potentials compared to that at open circuit during start-up and in the 4^th^ enrichment.

The activity of OTU1998 in the presence of an anode is consistent with our previous findings and may be a result of its interaction with electron-donating partners [51]. Meanwhile, the upregulation of its 16S rRNA expression by the cathodic potential can serve as evidence of OTU1998’s ability to accept extracellular electrons from the electrode.

*Geobacter*-related OTU1634 and OTU1635 had the highest transcript abundance in most of the samples (SI Figure S7). OTU1634 was particularly active under alternating polarity and open circuit, showing a transcript abundance of 9% at the end of the enrichment under those conditions (Figure 5A). This population responded to electrochemical stimulation in a similar manner to the methanogen. In the 4^th^ enrichment, anodic and cathodic potentials caused a 1.3-fold and 2.2-fold increase in transcript abundance, respectively, compared to open circuit (Figure 5B). The 16S rRNA expression of OTU1635 was also more active when the electrode potentials were applied but showed an opposite trend. Specifically, in the 4^th^ enrichment, the transcript abundance at the anodic potential was 3.2 folds relative to that at open circuit, whereas the change at the cathodic potential was only 0.9 folds (Figure 5B). The high transcript abundance of OTU1634 and OTU1635 demonstrates their electroactivity and importance in electro-methanogenic communities, but the different electrochemical responses are indicative of their different roles in the communities.

To further understand the interactions between the *Methanobacterium* and *Geobacter* populations, Bayesian networks were constructed with the core populations and three environmental factors (biogas production, pH, and cathodic current) that showed electrode potential-dependent variation. Bayesian networks are probabilistic graphical models capable of inferring the causal relationships between variables via directed acyclic graphs [86]. The networks inferred that OTU1998 could be affected by ten parent nodes (Figure 6A). Among them, OTU1634 had the highest positive network parameter (0.42) followed by OTU1066 (0.14) and OTU1635 (0.09). The cathodic current was inferred to exert a slightly negative impact on OTU1998 (network parameter -0.06). Using the actual values of the ten nodes, the abundance of OTU1998 was predicted with an R^2^ of 0.72 and an RMSE of 7.8% (Figure 6B). Considering that the prediction was performed at the OTU level, the accuracy was acceptable compared to previous studies [71, 87]. Null predictions were performed to reveal the influence of cathodic current, OTU1634, and OTU1635 on OTU1998 in a more quantitative manner. When the mean values of these nodes were fed into the networks, the predictions became noticeably less accurate (Figures 6C – 6E). Particularly, the R^2^ and RMSE of the null prediction with OTU1634 dropped to 0.30 and 12.4%, respectively, underpinning the close relationship between this *Geobacter* population and the methanogen.

**Figure 6.**
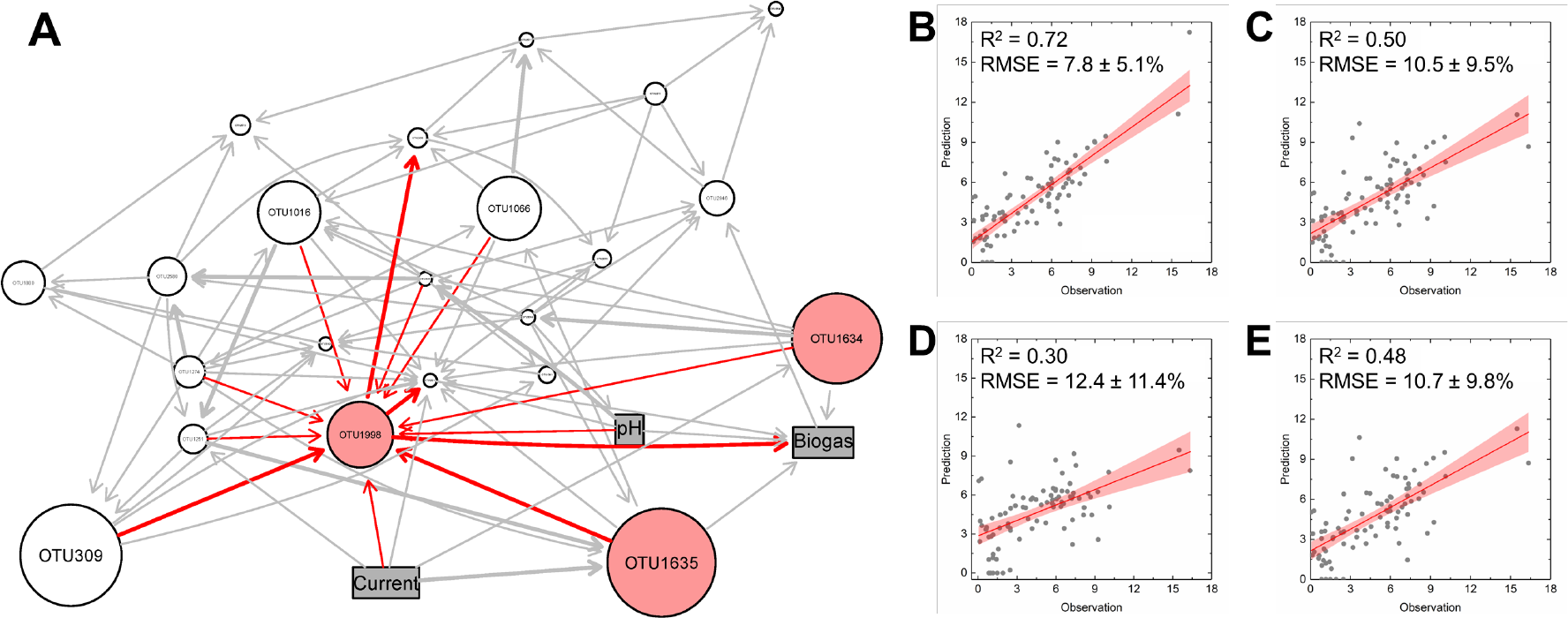
(A) Bayesian network constructed with the core populations (circle nodes) and three environmental factors (rectangular nodes). The size of the circle nodes represents the average abundance across all samples. Methanogens and *Geobacter* are highlighted in red. Prediction of the abundance of *Methanobacterium*-related OTU1998 was performed with (B) the actual values of all nodes, (C) the mean value of electric current and the actual values of all other nodes, (D) the mean abundance of OTU1634 and the actual values of all other nodes, (E) the mean abundance of OTU1635 and the actual values of all other nodes.

### 3.5 Perspectives

Electro-methanogenic microbial communities have attracted extensive interest due to their high efficiency in biogas production [17, 20, 42]. Significant efforts have been made to cultivate electro-methanogenic communities in engineered systems to improve biogas production [41, 43, 88]. The communities cultivated with conventional strategies, including cathode enrichment, microbial electrolysis, and conductive material amendment, are susceptible to perturbations [40, 44]. We attribute the unstable performance to the absence of electroactive bacteria, key electron-donating partners of electrotrophic methanogens [51]. Based on previous studies [54-56], we hypothesize that electroactive bacteria and electrotrophic methanogens can be simultaneously selected with alternating polarity (Figure 1), resulting in electro-methanogenic communities capable of robust biogas production. The findings of the present study collectively support our hypothesis. At the end of the cultivation, the cumulative biogas produced under alternating polarity was 11% higher than that under continuous cathode and 50% higher than that under open circuit (Figure 2). The communities cultivated with alternating polarity were structurally different from those under other conditions (Figure 3). The cultivation strategy may have important environmental implications. For example, the biogas production rate of existing full-scale systems can be readily enhanced by introducing cultivated activated carbon harboring electro-methanogenic communities (i.e., via bioaugmentation).

In addition to practical applications, electro-methanogenic communities cultivated with alternating polarity may serve as a model system to provide insight into the ecophysiology of key populations, particularly electrotrophic methanogens. The high abundance and activity of the putative electrotrophic methanogen OTU1998 at anodic and cathodic potentials (Figures 4 & 5, SI Figures S6 & S7) are in agreement with previous findings [22, 88-90], demonstrating the ability of some of the *Methanobacterium* spp. to accept electrons from both electrodes and electroactive bacteria. However, the metabolic pathways involved in extracellular electron uptake from the two electron donors remain elusive. In our prior work, we recovered the draft genomes of three anode-cultivated *Methanobacterium* populations with high quality (i.e., >98% completeness and <2% contamination). Multiple copies of the *mvhB* gene were found in their genomes and were more actively expressed under electrochemical stimulation. The *mvhB*-encoded iron-rich polyferredoxins were speculated to be responsible for intracellular electron transfer [91, 92]. It is possible that electrotrophic *Methanobacterium* use polyferredoxins to shuttle extracellular electrons to the heterodisulfide reductase/[NiFe]–hydrogenase complex for flavin-based electron bifurcation and methanogenesis [93, 94].

Although the electro-methanogenic communities cultivated in this study probably prefer to carry out electrotrophic methanogenesis, hydrogenotrophic methanogenesis may also affect biogas production. This is due to the fact that *Methanobacterium* can grow syntrophically with hydrogen-producing bacteria, e.g., *Ca*. G. *eutrophica*. Our previous studies revealed the ability of this novel *Geobacter* sp. to conserve energy through proton reduction [51, 82]. In the anodic cycle of alternating polarity, most of the extracellular electrons should have been channeled to the electrode (Coulombic efficiency 54%). However, biogas was still produced (Figure 2), implying the occurrence of hydrogenotrophic methanogenesis. Moreover, the inferred association between *Methanobacterium*-related OTU1998 and *Ca*. G. *eutrophica*-related OTU1634 (Figure 6D) may result from the involvement of both DIET and interspecies hydrogen transfer. Such metabolic versatility is, in principle, advantageous as it allows the two partners to cope with perturbations, thereby outcompeting other populations and dominating the community. The detailed ecophysiology of electro-methanogenic communities may help improve the cultivation strategy and thus warrants further investigation.

Although alternating polarity has been proven promising by this study, the electrode potential remains to be optimized in future studies. In particular, the anodic potential can have a strong impact on the community structure [95, 96]. In communities cultivated with anodes, *Geobacter* spp. were constantly observed with high abundance, but the community structure and microbial diversity could shift significantly as the anodic potential varied [97-100]. While the effect of anodic potential on electro-methanogenic communities is still inconclusive, it may be more favorable to apply lower anodic potentials (e.g., +0.2 V vs. SHE) in the case of alternating polarity. A moderate anodic potential allows some of the extracellular electrons to be channeled to electrotrophic methanogens during the selection of electroactive bacteria [51, 98], thereby maintaining the methanogenic activity of the communities.

The effect of electrode potential can be more complicated when alternation frequency is considered. Limited evidence suggests that high frequency does not favor the assembly of electro-methanogenic communities [58, 59]. For example, alternating the polarity every 10 minutes or 2 hours did not lead to the selection of methanogens or *Geobacter*, and the resulting communities did not produce biogas [60, 61]. It is possible that the alternation frequency should be longer than the doubling time of the populations to be selected so that they have sufficient time to adapt to the electrochemical stimulation [60, 61]. Moreover, the alternation frequency can be asymmetric, and a longer cathodic cycle may be more favorable, considering that methanogens typically have a longer doubling time than bacteria. The entangled effects of electrode potential and alternation frequency on community structure and microbial activity need to be better understood.

## Supporting information

Sumpplementary

## Supporting Information

Additional figures: schematics of the bioelectrochemical systems and cultivation procedure; methane content and COD removal in the 4^th^ enrichment; effluent pH under different cultivate conditions; current production under intermittent cathode and continuous cathode in the 4^th^ enrichment; first derivative of the voltammogram recorded at 0.2 mV/s and peak current vs. square root of scan rate; phylogenetic tree and relative abundance of the 22 core OTUs during start-up and in the four enrichments; activity (relative abundance of 16S rRNA transcript) of the 22 core populations under different cultivation conditions.

## Acknowledgment

This work was supported by the National Science Foundation (CBET-2128365) and the Global Energy Ecosystems (GE^2^) Research Development Program of the Office of Research, Innovation & Economic Development at the University of Tennessee, Knoxville (UTK).

## Reference

1. Appels, L., et al., Principles and potential of the anaerobic digestion of waste-activated sludge. Progress in Energy and Combustion Science, 2008. 34(6): p. 755–781.

2. Xu, F., et al., Anaerobic digestion of food waste – Challenges and opportunities. Bioresource Technology, 2018. 247: p. 1047–1058.

3. USDA, DOE, and USEPA, Biogas Opportunities Roadmap. 2014.

4. Batstone, D.J. and B. Virdis, The role of anaerobic digestion in the emerging energy economy. Current Opinion in Biotechnology, 2014. 27(0): p. 142–149.

5. Angelidaki, I., et al., Biogas upgrading and utilization: Current status and perspectives. Biotechnology Advances, 2018. 36(2): p. 452–466.

6. Schink, B. and A.J. Stams, Syntrophism among prokaryotes, in The prokaryotes. 2013, Springer. p. 471–493.

7. Campanaro, S., et al., Metagenomic analysis and functional characterization of the biogas microbiome using high throughput shotgun sequencing and a novel binning strategy. Biotechnology for Biofuels, 2016. 9(1): p. 1–17.

8. Nobu, M.K., et al., Microbial dark matter ecogenomics reveals complex synergistic networks in a methanogenic bioreactor. The ISME Journal, 2015. 9(8): p. 1710–1722.

9. Narihiro, T., et al., The nexus of syntrophy-associated microbiota in anaerobic digestion revealed by long-term enrichment and community survey. Environmental Microbiology, 2015. 17(5): p. 1707–1720.

10. Chen, Y., J.J. Cheng, and K.S. Creamer, Inhibition of anaerobic digestion process: A review. Bioresource Technology, 2008. 99(10): p. 4044–4064.

11. Stams, A.J. and C.M. Plugge, Electron transfer in syntrophic communities of anaerobic bacteria and archaea. Nature Reviews Microbiology, 2009. 7(8): p. 568–577.

12. Morris, B.E., et al., Microbial syntrophy: interaction for the common good. FEMS microbiology reviews, 2013. 37(3): p. 384–406.

13. Zhu, X., et al., Metabolic dependencies govern microbial syntrophies during methanogenesis in an anaerobic digestion ecosystem. Microbiome, 2020. 8(1): p. 22.

14. Lovley, D.R., Electrotrophy: Other microbial species, iron, and electrodes as electron donors for microbial respirations. Bioresource Technology, 2021:p. 126553.

15. Rotaru, A.-E., et al., Link between capacity for current production and syntrophic growth in Geobacter species. Frontiers in Microbiology, 2015. 6(744).

16. Gao, K. and Y. Lu, Putative extracellular electron transfer in methanogenic archaea. Frontiers in Microbiology, 2021. 12: p. 611739.

17. Cheng, S., et al., Direct Biological Conversion of Electrical Current into Methane by Electromethanogenesis. Environmental Science & Technology, 2009. 43(10): p. 3953–3958.

18. Walker, D.J., et al., The archaellum of Methanospirillum hungatei is electrically conductive. MBio, 2019. 10(2).

19. Holmes, D.E., et al., A membrane-bound cytochrome enables Methanosarcina acetivorans to conserve energy from extracellular electron transfer. MBio, 2019. 10(4).

20. Rotaru, A.-E., et al., A new model for electron flow during anaerobic digestion: direct interspecies electron transfer to Methanosaeta for the reduction of carbon dioxide to methane. Energy & Environmental Science, 2014. 7(1): p. 408–415.

21. Li, H., et al., Direct interspecies electron transfer accelerates syntrophic oxidation of butyrate in paddy soil enrichments. Environmental Microbiology, 2015. 17(5): p. 1533–1547.

22. Zheng, S., et al., Methanobacterium Capable of Direct Interspecies Electron Transfer. Environmental Science & Technology, 2020. 54(23): p. 15347–15354.

23. Lovley, D.R., Happy together: microbial communities that hook up to swap electrons. The ISME Journal, 2016. 11: p. 327.

24. Holmes, D.E., et al., Metatranscriptomic evidence for direct interspecies electron transfer between Geobacter and Methanothrix species in methanogenic rice paddy soils. Applied and Environmental Microbiology, 2017. 83(9): p. e00223–17.

25. Holmes, D.E., et al., Mechanisms for electron uptake by Methanosarcina acetivorans during direct interspecies electron transfer. MBio, 2021. 12(5): p. e02344–21.

26. Storck, T., B. Virdis, and D.J. Batstone, Modelling extracellular limitations for mediated versus direct interspecies electron transfer. The ISME Journal, 2015. 10: p. 621.

27. Zhao, Z., et al., Enhancing syntrophic metabolism in up-flow anaerobic sludge blanket reactors with conductive carbon materials. Bioresource Technology, 2015. 191: p. 140–145.

28. Lee, J.-Y., S.-H. Lee, and H.-D. Park, Enrichment of specific electro-active microorganisms and enhancement of methane production by adding granular activated carbon in anaerobic reactors. Bioresource Technology, 2016. 205: p. 205–212.

29. Chen, S., et al., Promoting interspecies electron transfer with biochar. Scientific Reports, 2014. 4: p. 5019.

30. Fu, Q., et al., Bioelectrochemical Analyses of the Development of a Thermophilic Biocathode Catalyzing Electromethanogenesis. Environmental Science & Technology, 2015. 49(2): p. 1225–1232.

31. Mei, R., et al., Novel Geobacter species and diverse methanogens contribute to enhanced methane production in media-added methanogenic reactors. Water Res, 2018. 147: p. 403–412.

32. Liu, Y., et al., Enhanced biogas production from swine manure anaerobic digestion via in-situ formed graphene in electromethanogenesis system. Chemical Engineering Journal, 2020. 389: p. 124510.

33. Liang, Y., A. Ma, and G. Zhuang, Construction of Environmental Synthetic Microbial Consortia: Based on Engineering and Ecological Principles. Frontiers in Microbiology, 2022. 13.

34. Shrestha, P.M. and A.-E. Rotaru, Plugging in or Going Wireless: Strategies for Interspecies Electron Transfer. Frontiers in Microbiology, 2014. 5.

35. Wang, L.-Y., et al., Expanding the Diet for DIET: Electron Donors Supporting Direct Interspecies Electron Transfer (DIET) in Defined Co-Cultures. Frontiers in Microbiology, 2016. 7(236).

36. Deutzmann, J.S. and A.M. Spormann, Enhanced microbial electrosynthesis by using defined co-cultures. The Isme Journal, 2016. 11: p. 704.

37. Mei, R. and W.-T. Liu, Quantifying the contribution of microbial immigration in engineered water systems. Microbiome, 2019. 7(1): p. 1–8.

38. Frigon, D. and G. Wells, Microbial immigration in wastewater treatment systems: analytical considerations and process implications. Current Opinion in Biotechnology, 2019. 57: p. 151–159.

39. Kim, J., et al., Ecogenomics-Based Mass Balance Model Reveals the Effects of Fermentation Conditions on Microbial Activity. Frontiers in Microbiology, 2020. 11(3115): p. 595036.

40. Bretschger, O., et al., Functional and taxonomic dynamics of an electricity-consuming methane-producing microbial community. Bioresource technology, 2015. 195: p. 254–264.

41. Wang, X.-T., et al., Enhancement of methane production from waste activated sludge using hybrid microbial electrolysis cells-anaerobic digestion (MEC-AD) process – A review. Bioresource Technology, 2022. 346: p. 126641.

42. Zakaria, B.S. and B.R. Dhar, Progress towards catalyzing electro-methanogenesis in anaerobic digestion process: Fundamentals, process optimization, design and scale-up considerations. Bioresource technology, 2019:p. 121738.

43. Martins, G., et al., Methane Production and Conductive Materials: A Critical Review. Environmental Science & Technology, 2018.

44. Ishii, S.i., et al., Bioelectrochemical stimulation of electromethanogenesis at a seawater-based subsurface aquifer in a natural gas field. Frontiers in Energy Research, 2019. 6: p. 144.

45. Logan, B.E., et al., Microbial fuel cells: methodology and technology. Environmental Science & Technology, 2006. 40(17): p. 5181–5192.

46. Zhao, L., et al., The underlying mechanism of enhanced methane production using microbial electrolysis cell assisted anaerobic digestion (MEC-AD) of proteins. Water Research, 2021. 201: p. 117325.

47. Lin, R., et al., Boosting biomethane yield and production rate with graphene: The potential of direct interspecies electron transfer in anaerobic digestion. Bioresource Technology, 2017. 239: p. 345–352.

48. Kato, S. and K. Igarashi, Enhancement of methanogenesis by electric syntrophy with biogenic iron‐sulfide minerals. MicrobiologyOpen, 2018:p. e00647.

49. Summers, Z.M., et al., Direct exchange of electrons within aggregates of an evolved syntrophic coculture of anaerobic bacteria. Science, 2010. 330(6009): p. 1413–1415.

50. Liu, X., et al., Syntrophic growth with direct interspecies electron transfer between pili-free Geobacter species. The ISME Journal, 2018.

51. Yuan, H., et al., Disentangling the syntrophic electron transfer mechanisms of Candidatus geobacter eutrophica through electrochemical stimulation and machine learning. Scientific Reports, 2021. 11(1): p. 15140.

52. Kawamata, Y., et al., Chemoselective electrosynthesis using rapid alternating polarity. Journal of the American Chemical Society, 2021. 143(40): p. 16580–16588.

53. Hayashi, K., et al., Chemoselective (Hetero) Arene Electroreduction Enabled by Rapid Alternating Polarity. Journal of the American Chemical Society, 2022. 144(13): p. 5762–5768.

54. Cheng, K.Y., G. Ho, and R. Cord-Ruwisch, Novel methanogenic rotatable bioelectrochemical system operated with polarity inversion. Environmental science & technology, 2011. 45(2): p. 796–802.

55. Mateos, R., et al., Impact of the start-up process on the microbial communities in biocathodes for electrosynthesis. Bioelectrochemistry, 2018. 121: p. 27–37.

56. Li, Z., et al., Polarity reversal facilitates the development of biocathodes in microbial electrosynthesis systems for biogas production. International Journal of Hydrogen Energy, 2019. 44(48): p. 26226–26236.

57. Wang, X., et al., Alternating current influences anaerobic electroactive biofilm activity. Environmental Science & Technology, 2016. 50(17): p. 9169–9176.

58. Yates, M.D., et al., Microbial electrochemical energy storage and recovery in a combined electrotrophic and electrogenic biofilm. Environmental Science & Technology Letters, 2017. 4(9): p. 374–379.

59. Riedl, S., et al., Cultivating electrochemically active biofilms at continuously changing electrode potentials. ChemElectroChem, 2019. 6(8): p. 2238–2247.

60. Izadi, P., et al., Bidirectional electroactive microbial biofilms and the role of biogenic sulfur in charge storage and release. Iscience, 2021. 24(8): p. 102822.

61. Mickol, R.L., et al., Metagenomic and Metatranscriptomic Characterization of a Microbial Community That Catalyzes Both Energy-Generating and Energy-Storing Electrode Reactions. Applied and Environmental Microbiology, 2021. 87(24): p. e01676–21.

62. Wang, B., et al., Intermittent electro field regulated mutualistic interspecies electron transfer away from the electrodes for bioenergy recovery from wastewater. Water Research, 2020. 185: p. 116238.

63. Narihiro, T., et al., Microbial Community Analysis of Anaerobic Reactors Treating Soft Drink Wastewater. PLOS ONE, 2015. 10(3): p. e0119131.

64. Isobe, K., et al., A simple and rapid GC/MS method for the simultaneous determination of gaseous metabolites. J Microbiol Methods, 2011. 84(1): p. 46–51.

65. Cao, L., et al., Comparative analysis of impact of human occupancy on indoor microbiomes. Frontiers of Environmental Science & Engineering, 2021. 15: p. 1–10.

66. Caporaso, J.G., et al., QIIME allows analysis of high-throughput community sequencing data. Nature Methods, 2010. 7(5): p. 335–336.

67. Callahan, B.J., et al., DADA2: High-resolution sample inference from Illumina amplicon data. Nature Methods, 2016. 13: p. 581.

68. Quast, C., et al., The SILVA ribosomal RNA gene database project: improved data processing and web-based tools. Nucleic Acids Research, 2013. 41(D1): p. D590–D596.

69. Bokulich, N.A., et al., Optimizing taxonomic classification of marker-gene amplicon sequences with QIIME 2’s q2-feature-classifier plugin. Microbiome, 2018. 6(1): p. 90.

70. Ludwig, W., et al., ARB: a software environment for sequence data. Nucleic Acids Research, 2004. 32(4): p. 1363–1371.

71. Yuan, H., et al., Unravelling and Reconstructing the Nexus of Salinity, Electricity, and Microbial Ecology for Bioelectrochemical Desalination. Environmental Science & Technology, 2017. 51(21): p. 12672–12682.

72. Scutari, M., Learning Bayesian Networks with the bnlearn R Package. Journal of Statistical Software. Journal of Statistical Software, 2010. 35(3): p. 1–22.

73. Bro, R., et al., Cross-validation of component models: a critical look at current methods. Analytical and bioanalytical chemistry, 2008. 390(5): p. 1241–1251.

74. He, Z., et al., Effect of electrolyte pH on the rate of the anodic and cathodic reactions in an air-cathode microbial fuel cell. Bioelectrochemistry, 2008. 74(1): p. 78–82.

75. Harnisch, F. and S. Freguia, A Basic Tutorial on Cyclic Voltammetry for the Investigation of Electroactive Microbial Biofilms. Chemistry – An Asian Journal, 2012. 7(3): p. 466–475.

76. Richter, H., et al., Cyclic voltammetry of biofilms of wild type and mutant Geobacter sulfurreducens on fuel cell anodes indicates possible roles of OmcB, OmcZ, type IV pili, and protons in extracellular electron transfer. Energy & Environmental Science, 2009. 2(5): p. 506–516.

77. Strycharz, S.M., et al., Application of cyclic voltammetry to investigate enhanced catalytic current generation by biofilm-modified anodes of Geobacter sulfurreducens strain DL1 vs. variant strain KN400. Energy & Environmental Science, 2011. 4(3): p. 896–913.

78. Bard, A.J. and L.R. Faulkner, Electrochemical methods: fundamentals and applications. Vol. 2. 1980:Wiley New York.

79. Ling, F., et al., Core-satellite populations and seasonality of water meter biofilms in a metropolitan drinking water distribution system. The ISME Journal, 2016. 10(3): p. 582–595.

80. Mayer, F., et al., Performance of different methanogenic species for the microbial electrosynthesis of methane from carbon dioxide. Bioresource technology, 2019. 289: p. 121706.

81. Poehlein, A., R. Daniel, and H. Seedorf, The draft genome of the non-host-associated methanobrevibacter arboriphilus strain dh1 encodes a large repertoire of adhesin-like proteins. Archaea, 2017. 2017.

82. Mei, R., et al., Novel Geobacter species and diverse methanogens contribute to enhanced methane production in media-added methanogenic reactors. Water Research, 2018. 147: p. 403–412.

83. Bond, D.R. and D.R. Lovley, Electricity Production by <em>Geobacter sulfurreducens</em> Attached to Electrodes. Applied and Environmental Microbiology, 2003. 69(3): p. 1548–1555.

84. Logan, B.E., et al., Electroactive microorganisms in bioelectrochemical systems. Nature Reviews Microbiology, 2019. 17(5): p. 307–319.

85. Biebl, H., Fermentation of glycerol by Clostridium pasteurianum—batch and continuous culture studies. Journal of Industrial Microbiology and Biotechnology, 2001. 27(1): p. 18–26.

86. Uusitalo, L., Advantages and challenges of Bayesian networks in environmental modelling. Ecological Modelling, 2007. 203(3): p. 312–318.

87. Kuang, J., et al., Predicting taxonomic and functional structure of microbial communities in acid mine drainage. The ISME Journal, 2016. 10(6): p. 1527–1539.

88. Gao, K., et al., Response of Methanogen Communities to the Elevation of Cathode Potentials in Bioelectrochemical Reactors Amended with Magnetite. Applied and Environmental Microbiology, 2021. 87(21): p. e01488–21.

89. Perona-Vico, E., et al., [NiFe]-hydrogenases are constitutively expressed in an enriched Methanobacterium sp. population during electromethanogenesis. PloS one, 2019. 14(4).

90. Baek, G., P.E. Saikaly, and B.E. Logan, Addition of a carbon fiber brush improves anaerobic digestion compared to external voltage application. Water Research, 2021. 188: p. 116575.

91. Hedderich, R., et al., Isolation and characterization of polyferredoxin from Methanobacterium thermoautotrophicum The mvhb gene product of the methylviologen-reducing hydrogenase operon. FEBS Letters, 1992. 298(1): p. 65–68.

92. Ferry, J.G., Methanogenesis: ecology, physiology, biochemistry & genetics. 2012:Springer Science & Business Media.

93. Kaster, A.-K., et al., Coupling of ferredoxin and heterodisulfide reduction via electron bifurcation in hydrogenotrophic methanogenic archaea. Proceedings of the National Academy of Sciences of the United States of America, 2011. 108(7): p. 2981–2986.

94. Watanabe, T. and S. Shima, MvhB-type Polyferredoxin as an Electron-transfer Chain in Putative Redox-enzyme Complexes. Chemistry Letters, 2021. 50(2): p. 353–360.

95. Torres, C.s.I., et al., Selecting Anode-Respiring Bacteria Based on Anode Potential: Phylogenetic, Electrochemical, and Microscopic Characterization. Environmental Science & Technology, 2009. 43(24): p. 9519–9524.

96. Zhu, X., et al., Microbial Community Composition Is Unaffected by Anode Potential. Environmental Science & Technology, 2014. 48(2): p. 1352–1358.

97. Ishii, S.i., et al., Microbial population and functional dynamics associated with surface potential and carbon metabolism. The ISME Journal, 2014. 8(5): p. 963–978.

98. Dennis, P.G., et al., Anode potential influences the structure and function of anodic electrode and electrolyte-associated microbiomes. Scientific reports, 2016. 6: p. 39114.

99. Hari, A.R., et al., Set anode potentials affect the electron fluxes and microbial community structure in propionate-fed microbial electrolysis cells. Scientific reports, 2016. 6(1): p. 1–11.

100. Ying, X., et al., The impact of electron donors and anode potentials on the anode-respiring bacteria community. Applied microbiology and biotechnology, 2017. 101(21): p. 7997–8005.

