## Supplementary material for "Alternating Polarity as a Novel Strategy for Cultivating Electro-Methanogenic Microbial Communities Capable of Robust Biogas Production": Sumpplementary

Type of contribution***: Research Article***

[*] Corresponding authors.
Heyang Yuan:
Qiang He:

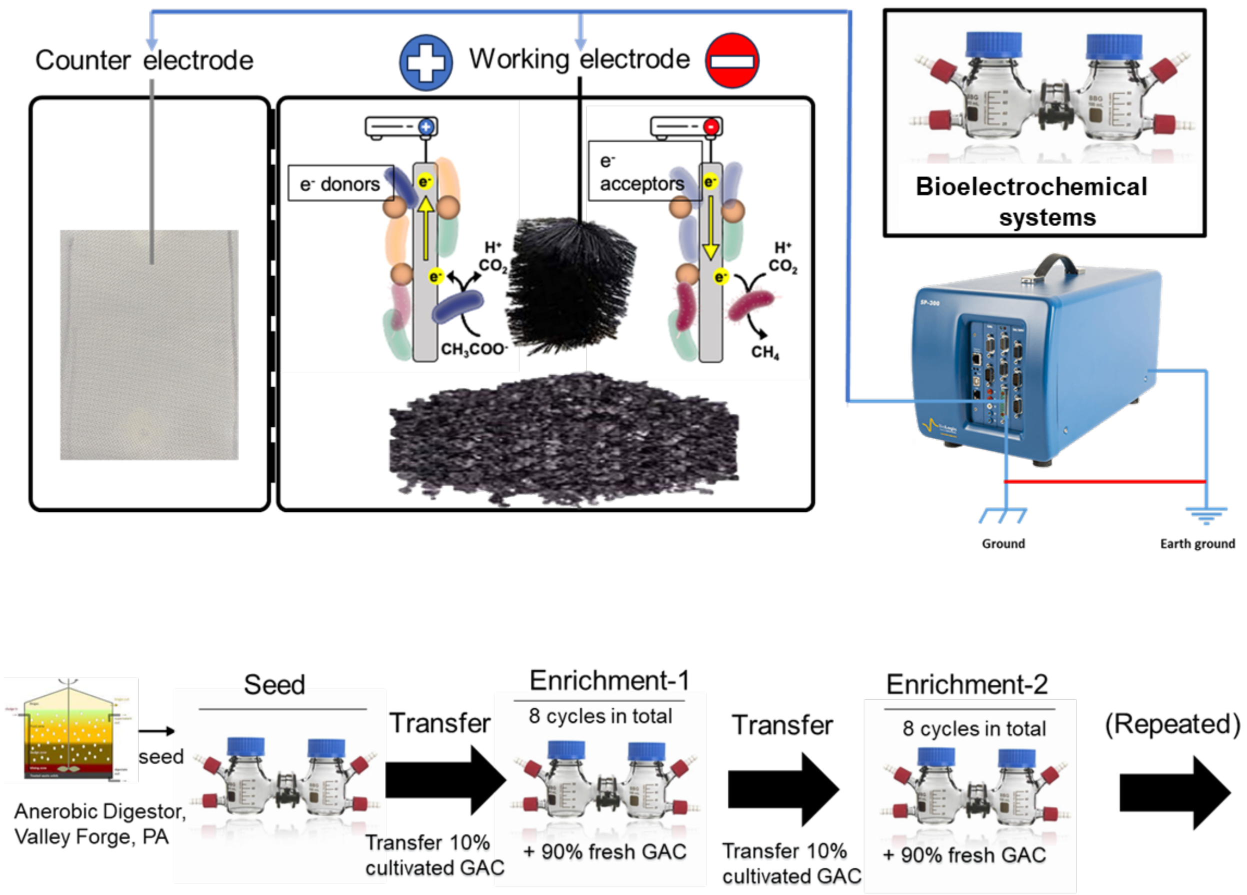


Figure S1. Schematics of the bioelectrochemical systems and cultivation procedure.

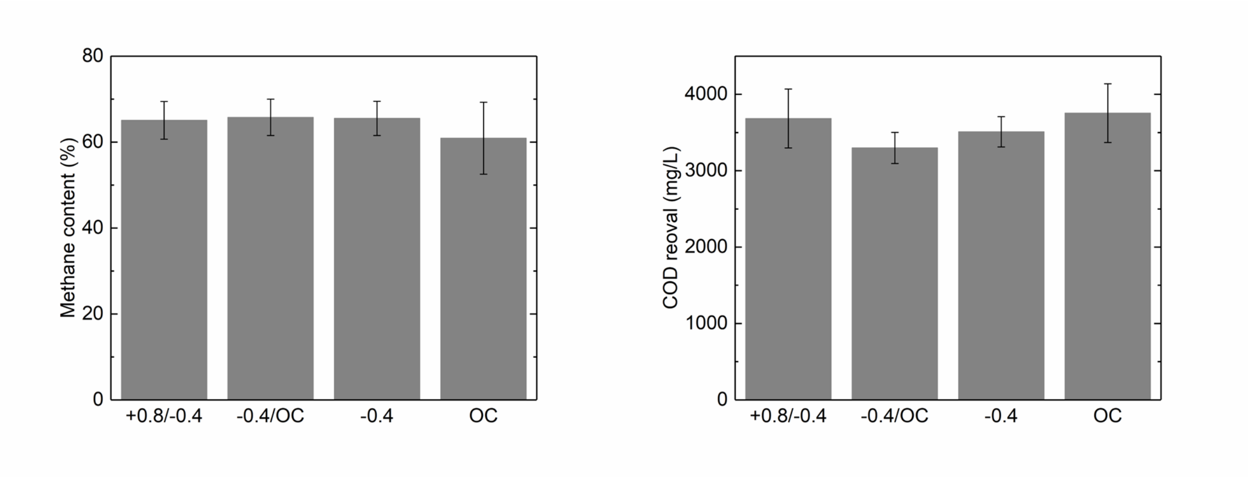


Figure S2. Methane content (left panel) and COD removal (right panel) in the 4$^{th}$ enrichment.

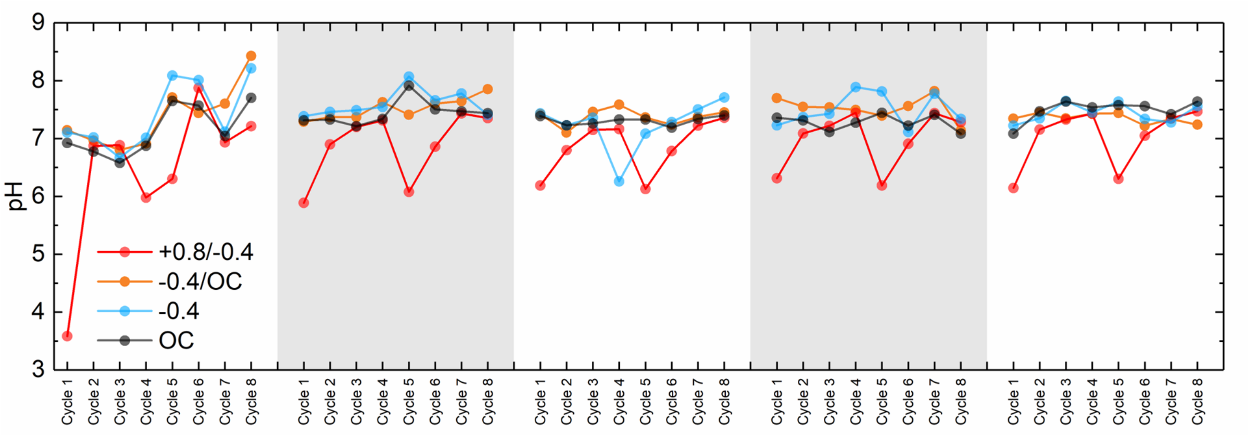

Figure S3. Effluent pH under different cultivate conditions.

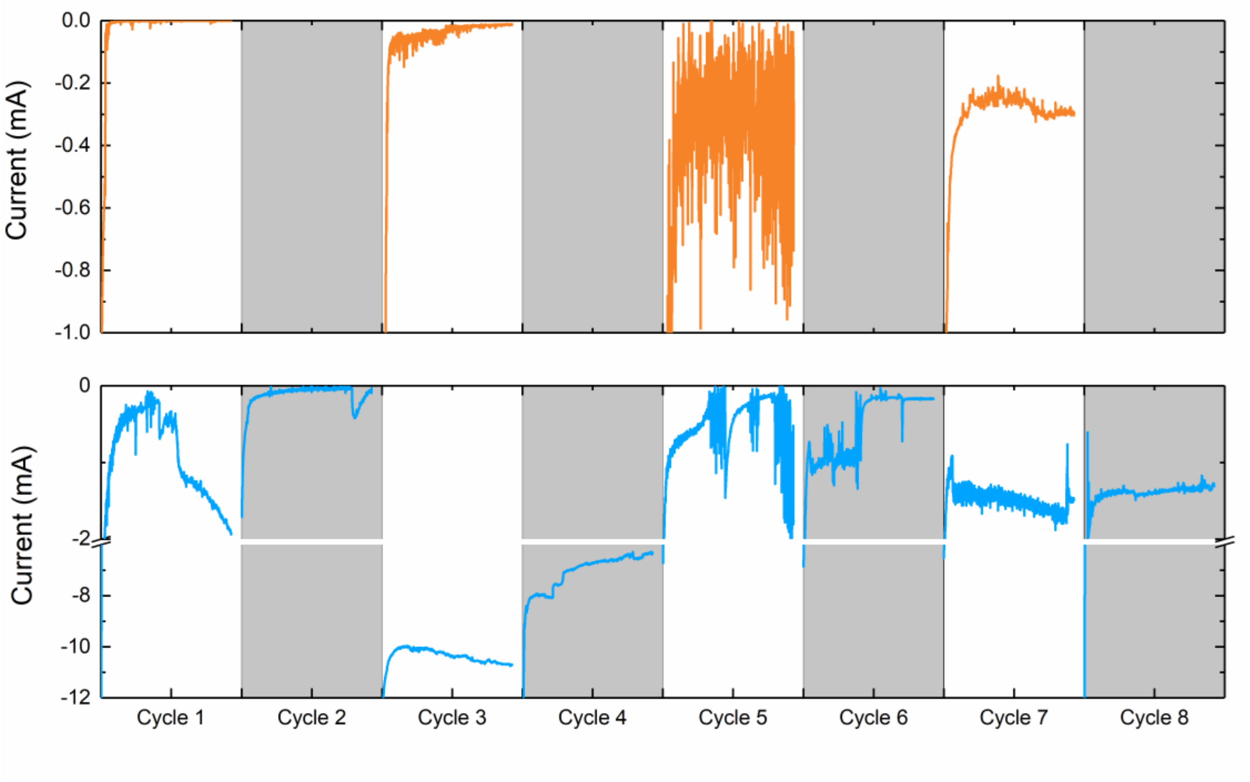


Figure S4. Current production under intermittent cathode (upper panel) and continuous cathode (lower panel) in the 4th enrichment.

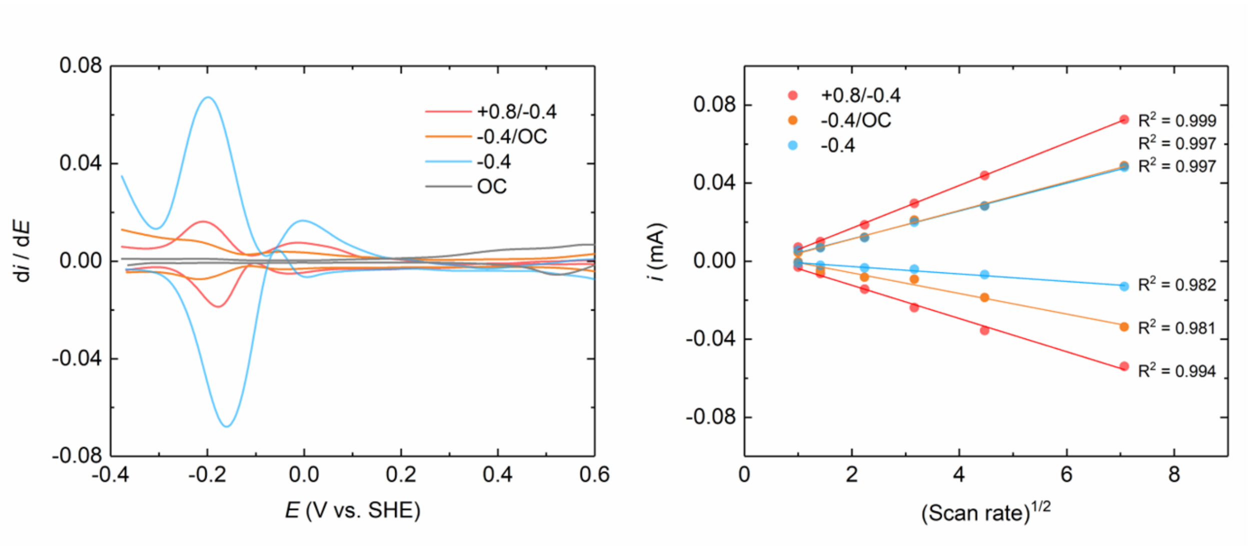


Figure S5. First derivative of the voltammogram recorded at 0.2 mV/s (left panel) and peak current vs. square root of scan rate (right panel).

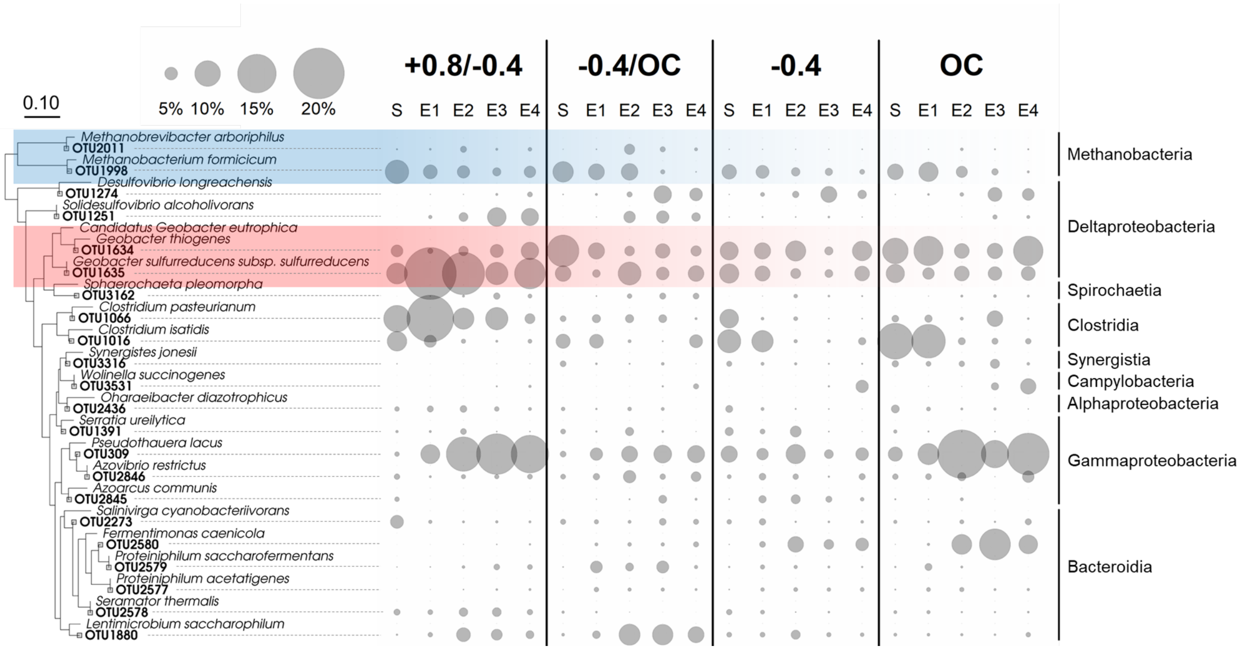


Figure S6. Phylogenetic tree and relative abundance of the 22 core OTUs during start-up and in the four enrichments.

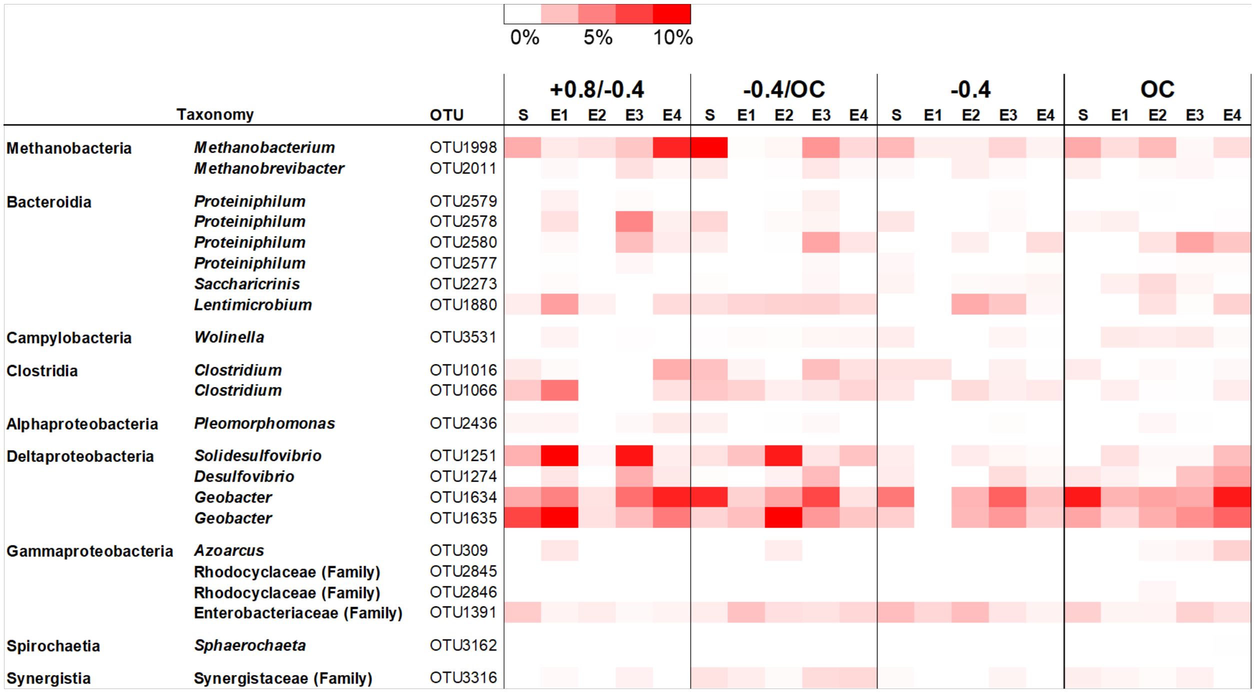


Figure S7. Activity (relative abundance of 16S rRNA transcript) of the 22 core populations under different cultivation conditions.
